# Variant calling using NGS and sequence capture data for population and evolutionary genomic inferences in Norway Spruce (*Picea abies*)

**DOI:** 10.1101/805994

**Authors:** Carolina Bernhardsson, Xi Wang, Helena Eklöf, Pär K. Ingvarsson

**Author notes:** These authors contributed equally (Carolina Bernhardsson, Xi Wang). Email addresses: CB, XW, HE, PKI.

## Abstract

Advances in next-generation sequencing methods and the development of new statistical and computational methods have opened up possibilities made for large-scale, high quality genotyping in most organisms. Conifer genomes are large and are known to contain a high fraction of repetitive elements and this complex genome structure has bearings for approaches that aim to use next-generation sequencing methods for genotyping. In this chapter we provide a detailed description of a workflow for variant calling using next-generation sequencing in Norway spruce (*Picea abies*). The workflow that starts with raw sequencing reads and proceeds through read mapping to variant calling and variant filtering. We illustrate the pipeline using data derived from both whole-genome resequencing data and reduced-representation sequencing. We highlight possible problems and pitfalls of using next-generation sequencing data for genotyping stemming from the complex genome structure of conifers and how those issues can be mitigated or eliminated.

## 1. Introduction

Conifers were one of the last plant groups lacking genome assemblies, but recently several draft genomes have become available for a number of conifers such as Norway spruce (*Picea abies*, Nystedt et al. 2013), Loblolly pine (*Pinus taeda*, Zimin et al 2014, 2017), Sugar pine (Stevens et al 2016) and Douglas Fir (Neale et al 2017). This has opened up new possibilities to assess genome-wide levels of genetic diversity in conifers. Earlier studies of genetic diversity in Norway spruce has either been limited to coding regions (e.g Huertz et al 2006, Chen et al. 2012) or have used various complexity reduction methods, such as genotyping by sequencing, restriction site associated sequencing, or targeted capture sequencing (Baison et al 2018) to estimate levels of genetic diversity within species. While we have learned a lot about levels of genetic diversity in Norway spruce from such studies, we still lack detailed information on, for instance, levels of nucleotide polymorphism and linkage disequilibrium in non-genic regions. However, with the availability of a reference genome sequences (Nystedt et al 2013), whole genome re-sequencing is now also possible in conifers such as Norway spruce.

Conifer genomes are large (20-40Gb) and have high repetitive content and current draft genome assemblies in conifers are therefore often fragmented into many, relatively short scaffolds. In addition, large fractions of the predicted genome sizes are also missing from reference genomes. The fragmented nature of conifer reference genome assemblies, combined with the high repetitive content make variant calling in conifers difficult. This is true regardless of what techniques have been used to generate sequencing data but perhaps more so for whole-genome re-sequencing data that can be expected to provide a relatively unbiased coverage of the target genome. In this chapter we review methods available for variant calling using NGS data and outline some of the issues one may face when performing analyses of data from whole-genome re-sequencing (WGS) in Norway spruce. In particular we discuss the performance of variant calling across different genomic contexts, such as coding and non-coding regions and regions known to be composed of repetitive elements. We also compare variant calling using WGS data with data derived from sequence capture probes, designed to target non-repetitive sequences in the *P. abies* genome and discuss how collapsed genomic regions in the assembly complicates the task of filtering for good reliable variant- and genotype calls. Having access to robust variant calls is important for downstream analyses, such as population genomic analyses or inferences of the demographic history of individuals, populations or the species as a whole. To highlight these issues, we end by assessing how different approaches to variant calling alter the site frequency spectrum of variants and hence possible evolutionary inferences drawn from the data.

## 2. Sample collection

We sampled 35 individuals of Norway spruce (*Picea abies*) spanning their natural distributions, mainly from Russia, Finland, Sweden, Norway, Belarus, Poland and Romania for use in whole genome re-sequencing. Individuals Pab001-Pab015 were all derived from unique populations and no specific measurements were taken when they were collected. Samples were taken from newly emerged needles or dormant buds for each individual and stored in −80 °C until DNA extraction. In contrast, individuals Pab016-Pab035 were sampled from two different areas, one in the eastern and one in the western part of Västerbotten province in northern Sweden. Two different populations were sampled in each area, one old and untouched forest (>100 years old) and one young planted population (<20 years old). For every population a transect was made and five trees were sampled from each population along the transect. From each tree, a number of fresh shoots was broken off and put into pre-labeled zip lock bags before being taken back to the lab for DNA extractions.

Genomic DNA was extracted using Qiagen plant mini kit following manufacturer’s instructions. All sequencing was performed at the National Genomics Initiative platform (NGI) at the SciLifeLab facilities in Stockholm, Sweden, paired-end libraries with an insert size of 500bp on different Illumina HiSeq platforms (Pab001-Pab006 on HiSeq 2000, remaining individuals on HiSeq X). The original location, estimated coverage from raw sequencing reads and coverage of mapped reads for each individual are given in Table 1.

Samples analysed using sequence capturing methods were obtained from Bernhardsson et al. (2018, 1997 haploid samples) and Baison et al. (2018, 526 diploid samples). For further information on sample collection, DNA extraction and sequencing please refer to the original publications.

## 3. SNP and genotype calling pipeline

Over the recent decades, Next-Generation Sequencing (NGS) technologies have developed at an extraordinary rapid pace and have led to ever decreasing costs per mega-base sequence generated. This has in turn led to rapid increase in the number and diversity of sequenced genomes (Goodwin et al. 2016). These new sequencing technologies have already facilitated and improved many research avenues in biology, such as transcriptomics, gene annotation and RNA splice identification (Bao et al. 2011). Moreover, meta-genomic (Schuster 2007) and genome methylation analysis (Morozova and Marra 2008) have also benefited from NGS technologies. However, NGS platforms provide higher associated error rates and generally shorter read lengths compared with traditional Sanger sequencing platforms and therefore require much more careful examinations of the results, particularly for variant discovery and downstream applications (Liu et al. 2012; Goodwin et al. 2016). Thus, meaningful analysis of NGS data relies crucially on accurate calling of SNPs and genotypes (Nielsen et al. 2011). Here we first highlight some of the issues that may contribute to ambiguities in SNP and genotype calling, e.g. repetitive DNA, or misalignment of sequence reads, and then review some recent algorithms and tools that are capable of improving the sensitivity and specificity of SNP detection and genotype determination from NGS data. The pipeline for SNP and genotype calling applied in this chapter includes initial reads mapping, post-alignment processing, pre-calling processing, final SNP and genotype calling and subsequent variant filtering (Figure 1). The steps of this pipeline are exemplified by using whole-genome re-sequencing data from 35 Norway spruce (*Picea abies*) individuals generated by different Illumina HiSeq platforms.

**Figure 1:**
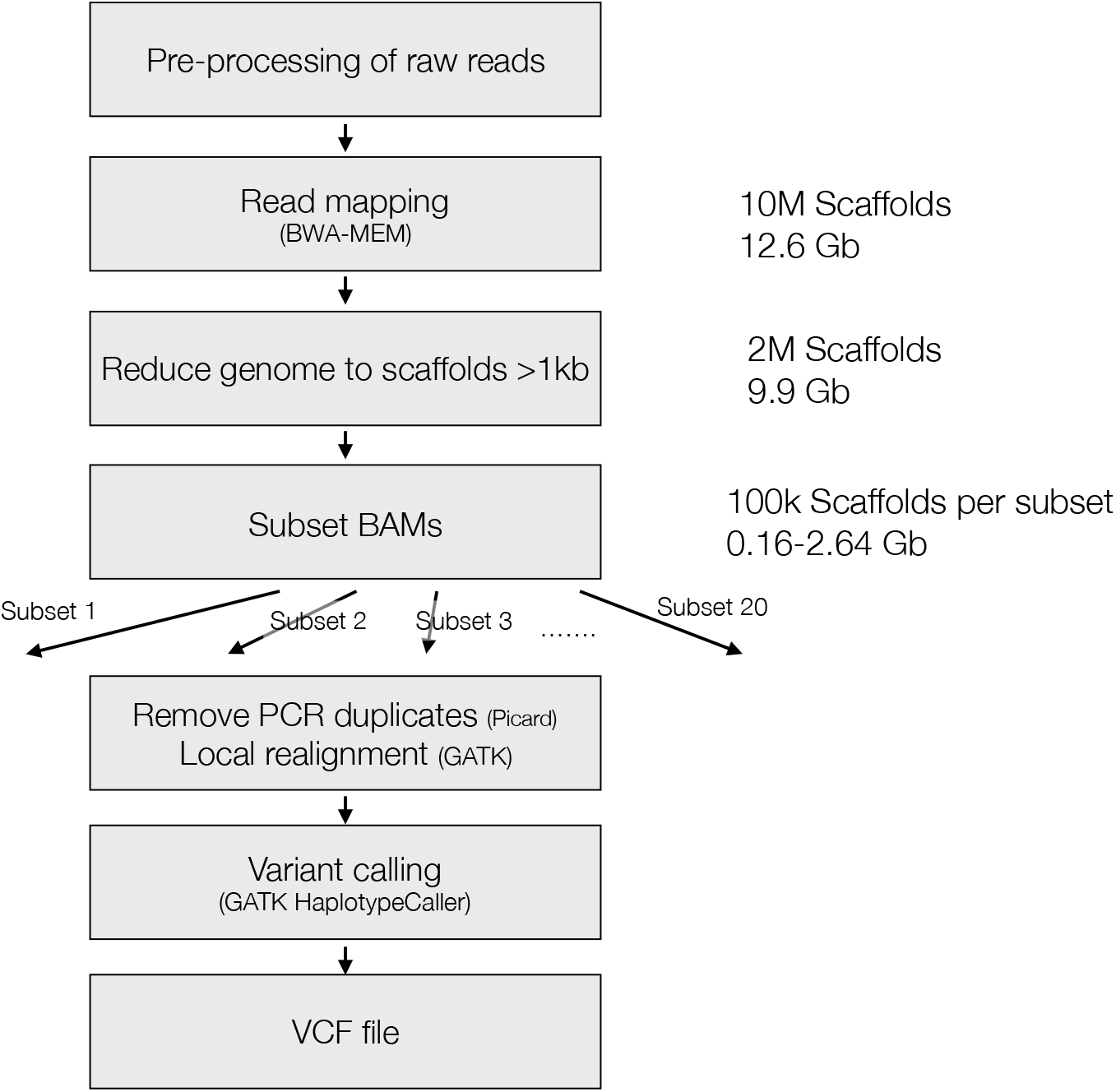
Pipeline of SNP and genotype calling based on Norway spruce. As a first step, paired end reads were mapped to the whole genome assembly of Norway spruce (~10 million scaffolds covering 12.6 Gb out of the ~20 Gb estimated genome size) using BWA-MEM with default settings. Sample BAM files were then reduced to only include scaffolds longer than 1 Kb (~2 million scaffolds covering 9.5 Gb of the genome), merged into a single BAM file per sample and then subdivided into 20 genomic subsets with ~100 K scaffolds in each. The genomic subset BAM files were then marked for PCR duplicates using `Picard`, subjected to local realignment around indels using `GATK RealignerTargetCreator` and `IndelRealigner` and genomic intermediate variant calling using `GATK Haplotypecaller` in gvcf mode. A joint genotype call over all 35 samples per genomic subset was then performed using `GATK GenotypeGVCFs` to ashieve the final raw vcf files (one file for each of the 20 genomic subset).

### 3.1 Initial reads mapping (step 1)

Mapping is the most fundamental step in variant detection using NGS and involves aligning raw or pre-processed sequence reads to a reference genome of either the same or a closely related species to reveal the location(s) of reads within the reference genome (Trapnell and Salzberg 2009; Flicek and Birney 2010; Wang et al. 2015). The accuracy of mapping plays a crucial role in variants calling (Nielsen et al. 2011) as incorrectly aligned reads may lead to errors in SNP and genotype calling, and therefore will restrict the accuracy of all downstream analyses. Although progressively more advanced NGS technologies feature several advantages such as high throughput, lower cost per base and higher accuracy when compared with traditional Sanger-based sequencers (Bao et al. 2011), several obstacles stemming from the inherent characteristics of NGS data need to be considered when performing this step.

#### 3.1.1 Short-read mapping

The first challenge is that reads from NGS platforms are usually short (most delivered read lengths fall in the range of 35 - 400 bp), when compared with the reads from Sanger sequencers (600 - 800 bp) (Bao et al. 2011; Mardis 2017). In order to use these short reads efficiently, the initial process, before actual alignment, involves the construction of indices for the reference sequence and/or for the raw short-read sequence data known as ‘indexing’, is intended to speed up subsequent mapping steps. The two most commonly used indexing algorithms have been incorporated in a large number of mapping software tools: (i) indexing algorithms based on hash tables, and (ii) indexing algorithms based on Burrows–Wheeler transform (BWT). A ‘Hash table’ is a common data structure which makes it possible to search rapidly through complex and non-sequential data using an index. One option is to scan the hash table of input reads using the reference genome. However, scanning a hash table of input reads usually requires smaller and more variable memory, while scanning the entire reference genome may use more processing time especially when there are relatively few reads in the input set (Flicek and Birney 2010). Another option is to scan the hash table of the reference genome by using the set of input reads. Regardless of the size of input reads, the memory of the reference hash table remains constant, but may on the other hand be large, depending on the size and complexity of the reference genome (Flicek and Birney 2010). Several software tools have been developed based on hash table algorithms, such as `MAQ` (Li et al. 2008a), `SOAP` (Li et al. 2008b), `SHRiMP` (Rumble et al. 2009) and `Stampy` (Lunter and Goodson 2011). BWT-based implementations are much faster compared to hash table based algorithm at the same sensitivity level (Flicek and Birney 2010), they are also more memory-efficient and are par-ticularly useful for aligning repetitive reads (Nielsen et al. 2011). Another advantage of the BWT approach is the ability to store the complete reference genome index on disk and load it completely into memory on almost all standard bioinformatics computing clusters (Flicek 2009; Flicek and Birney 2010). Currently, the most widely used software tools based on the BWT algorithms include `Bowtie` (http://bowtie.cbcb.umd.edu/; Langmead et al. 2009), the `Burrows–Wheeler Aligner` (`BWA`) (http://maq.sourceforge.net/bwa-man.shtml; Li and Durbin 2009), and `SOAP2` (http://soap.genomics.org.cn/; Li et al. 2009a).

#### 3.1.2 Extremely large amounts of data

Another challenge for working with NGS data is that the amount produced by most NGS methods are orders of magnitude greater than that generated by earlier techniques (Flicek and Birney 2010). For example, Illumina (HiSeq 2000) and Applied Biosystems (ABI SOLiD 4) can deliver hundreds of millions of sequences per run, while Sanger-style sequencer only produces thousands of reads per run (Hung and Weng 2017). Therefore, in order to map a vastly greater numbers of reads (millions or billions) to a reference genome, any algorithm must be optimized for speed and memory usage, especially when reference genomes are very huge. Accordingly, several very memory efficient short-read alignment programs have been developed. For example, `Bowtie` (http://bowtie-bio.sourceforge.net/index.shtml; Langmead et al. 2009) and `BWA`, as examples of the two most efficient short-read aligners, achieve throughputs of 10–40 million reads per hour on a single computer processor (Treangen and Salzberg 2012).

#### 3.1.3 Repetitive DNA mapping

Repetitive DNA sequences, which are similar or identical to sequences elsewhere in the genome, are abundant across a broad range of organisms, from bacteria to mammals (Treangen and Salzberg 2012). Mapping repetitive DNA sequences to a reference genome create ambiguities of deciding what to do with reads that map to multiple locations, which, in turn, can produce errors when interpreting results. Mapping of repetitive sequencing reads is therefore one of the most commonly encountered problem in read mapping. One possible solution to this problem is to simply remove all multiple mapped reads. However, this discarding strategy may complicate the calculation of coverage by reducing coverage in an uneven fashion (Li et al. 2008a), and may result in many undetected biologically important variants when analyses are performed across unique regions involved in the repetitive contents in the genome. Some repeats have already proved to play important roles in, for instance, human evolution (Jurka et al. 2007; Britten 2010), sometimes creating novel functions while sometimes acting as independent ‘selfish’ genetic elements (Kim et al. 2008; Hua-Van et al. 2011; Treangen and Salzberg 2012). An alternative option is to only report the region with the fewest mismatches (e.g. as done in `BWA` and `Bowtie`). Specifically, when there are multiple equally good best alignment matches, the aligner can either randomly choose one or choose to report all such alignments (e.g. `SOAP2`). Among those aligners reporting all matches, another choice is to report up to a maximum number (`m`) by ignoring multiple reads that align to `>m` locations. To deal with these issues, the concept of ‘mapping quality’ was introduced in several software tools to enable evaluation of the likelihood of correct mapping of reads by considering a number of factors, such as base qualities, the number of base mismatches and/or the existence and size of gaps in the alignment (Li et al. 2008a; Wang et al. 2015). For example, in order to evaluate the reliability of alignments, `MAQ` assigns a Phred-scaled quality score which measures the probability that the true alignment is not the one found by `MAQ` for each individual alignment (Li et al. 2008a). However, `MAQ`’s formula overestimates the probability of missing the true hit, which results in an underestimation of mapping quality (Li and Durbin 2009). `BWA` was developed with a similar algorithm as `MAQ` but has been modified by assuming the true hit can always be found. Simulation reveals that BWA may therefore overestimate mapping quality, although the deviation is relatively small (Li and Durbin 2009).

In general, the choice of alignment tool and the corresponding parameter settings are very important because the outcome will significantly influence the accuracy of variant calling and further downstream analysis. `BWA`, which is based on the BWT algorithm, is much faster compared to many other programs based on hash table algorithm at the same sensitivity level, has more efficiently memory usage, which is par-ticularly useful for aligning repetitive reads, and has a smaller deviation of mapping quality. This is therefore often the best choice for mapping raw sequence reads to a reference genome. `BWA` provides three different mapping methods: 1) `BWA-MEM` that is usually suggested for 70bp or longer Illumina, 454, Ion Torrent and Sanger reads, and is also generally recommended for high-quality queries as it is faster and more accurate; 2) `BWA-backtrack` is especially suited for short sequences and 3) `BWA-SW` which may have better sensitivity when alignment gaps are frequent. Consequently, in our whole genome re-sequencing project of Norway spruce, we have employed `BWA-MEM` to align all paired-end reads to the reference genome.

### 3.2 Post-alignment processing (step 2)

After mapping reads to the reference, various post-alignment data processing are usually suggested to facilitate further analytical steps. The most common tasks include output file manipulation (e.g. format converting, indexing) and the creation of summary reports from the alignment process. Appropriate formats of output from mapping not only ensure downstream compatibility with variant callers, such as ‘Haplotypecaller’ from the Genome Analysis Toolkit package (`GATK`) (McKenna et al. 2010), but also allow for a reduction of the output data set size, for example, transferring from SAM (a text-based format for storing biological sequences aligned to a reference sequence which is developed by Li et al. 2009c) file to the binary version (BAM) file (Altmann et al. 2012; Mielczarek and Szyda 2016). The SAMtools package can be used to manipulate a variety of functions on output files, such as, format changing, sorting, indexing, or merging (Li et al. 2009c). Moreover, a detailed summary statistics of aligned output are necessary for evaluating the overall quality and correctness of alignment, which benefits for following steps such as SNP detection. Some aligners usually generate a simple summary describing the alignment process (e.g. `Bowtie2, SOAP2`), while some are not available, such as `BWA`. Thus, it is possible to generate a summary report by using other software, e.g. the `SAMtools flagstat` (Li et al 2009c) and `CollectAlignmentSummaryMetric`s module of `Picard` (http://broadinstitute.github.io/picard/).

After mapping raw sequencing reads to the reference genome using `BWA-MEM`, we have employed `SAMtools` to manipulate output files, for instance for sorting and indexing. Besides, we also used `flagstat` in `SAMtools` to generate summary statistic reports including e.g. the total number of reads that pass or fail QC (quality controls), in order to evaluate the quality of the alignments produced during read mapping. In addition, considering the characteristics of Norway spruce genome which has an extremely large genome size (~20Gb) and very high repetitive content (~70%) (Nystedt et al. 2013), it is challenging to accurately call variants both from a computational perspective and from the large data volumes that need to be handled and stored in this project. Before SNP and genotype calling, we therefore performed several steps to reduce the computational complexity of the SNP calling process. First we split both the output file from the mapping step and the reference into smaller subsets so that multiple data sets can be manipulated in parallel. Also, smaller data sets (in this case number of scaffolds) are crucial to allow software tools, such as GATK, to function properly.

We first reduced the reference genome by only keeping genomic scaffolds greater than 1kb in length using `bioawk` (https://github.com/lh3/bioawk). All BAM files were then subsetted using the reduced reference genome with the `view` module in `SAMtools`. All reduced BAM files for each individual were merged into a single BAM file using the `merge` module in `SAMtools` (Li et al. 2009c). The final step was to split the merged BAM files of each individual into 20 genomic subsets, with each subset containing roughly 100,000 scaffolds, using `SAMtools view`. We simultaneously subdivided the reference genome into the corresponding 20 genomic subsets by keeping exactly same scaffolds by using `bedtools` (Quinlan and Hall 2010). Indexing was performed for both BAM subsets and reference subsets to make them available for subsequent analyses.

### 3.3 Pre-calling processing (step 3)

Additional processing steps are usually recommended before proceeding to the actual variant calling, so that variant detection results can be more reliable (Li et al. 2009c; McKenna et al. 2010; Altmann et al. 2012; Mielczarek and Szyda 2016).

#### 3.3.1 PCR duplicates

DNA amplification by polymerase chain reaction (PCR) has been a necessary step in the preparation of libraries for most of the NGS platforms, such as Illumina, 454, IonTorrent and SOLiD (Mardis 2008; Morozova and Marra 2008; Escalona et al. 2016). The number of duplicate clones in the library will increase if there are too many PCR cycles or when obtaining DNA from a gel slice to get uniform fragment lengths (Li et al. 2009c). Consequently, the same read pairs may occur many times in the raw data and this will result in unexpectedly high depths in some regions, resulting in a skewed coverage distribution (Mielczarek and Szyda2016). Further, excessively high read depth could potentially lead to large frequency differences between the two alleles of a heterozygous site (Li et al. 2009c), which may subsequently bias the number of variants and may substantially influence the accuracy of the variant detection (Wang et al. 2005). Removal of these artifacts is thus an essential pre-calling processing step, in particular on applications based on re-sequencing data. `Picard MarkDuplicates` (http://broadinstitute.github.io/picard/) and `Samtools Rmdup` (Li et al. 2009c) are two commonly used software which allow the user to either mark these duplicate reads or completely remove them from the alignment. Both methods rely on similar approaches, however, compared with `rmdup, MarkDuplicates` is likely a better choice as it also handles inter-chromosomal read pairs, considers the library for each read pair, keeps a read pair from each library and does not actually remove reads but rather flags the duplicates by setting the SAM flag to 1024 for all but the best read pair (Ebbert et al. 2016).

#### 3.3.2 Local realignment

Another important preprocessing step involves alignment corrections. Artefacts in alignment, usually occurring in regions with insertions and/or deletions (indels) during the mapping step, can result in many mismatching bases relative to the reference in regions of misalignment. Such misaligned regions can be easily mistaken as SNPs during variant calling. Read mapping proceeds by independently mapping each read, and reads covering an indel at the start or end of the read can often be incorrectly mapped even when other reads are correctly mapped across the indel. To alleviate these issues, local realignment is usually performed to realign reads in the vicinity of an identified misalignment in order to minimise the number of mismatching bases (Mielczarek and Szyda 2016). Many software tools have been developed to perform local realignment, such as `SRMA` (Homer and Nelson 2010), which realign reads only in color space originating from the `SOLiD` platform. `GATK` (McKenna et al. 2010) is probably the most commonly used software for performing local realignment and it proceeds in two steps: 1) determining (small) suspicious intervals which are likely in need of realignment (`RealignerTargetCreator`), and 2) running the actual realigner over those intervals (`IndelRealigner`).

In this project, we first marked potentially PCR duplicates by using `MarkDuplicates` in `Picard`. We then performed local realignment by first detecting suspicious intervals using `RealignerTargetCreator`, followed by realignment of those intervals using `IndelRealigner`, both implemented in `GATK` (DePristo et al. 2011). Both of these steps of pre-call processing were run separately on the 20 genomic subsets of each individual (700 subsets in total across all 35 individuals).

### 3.4 SNP and genotype calling (step 4)

The development of NGS has made it possible to identify a large number of variants in almost any organism. Sequencing entire genomes potentially allows for the discovery of all existing polymorphisms, and can thus identify not only common but also rare SNPs which have been implicated in controlling many complex phenotypes (Mielczarek and Szyda 2016). This means that it should be possible to, at least in theory, identify the true causal mutations directly from resequencing data rather than relying on using linkage disequilibrium between unknown causal mutations and marker SNPs. However, to achieve this, SNP detection and genotype calling need to be performed across polymorphic sites in the NGS data. SNP calling, also known as variant calling, is aimed at detecting sites which differ from a reference sequence, while genotype calling is the process of determining the actual genotype for each individual based on positions in which a SNP or a variant has already been called (Nielsen et al. 2011; Li et al. 2013). SNP calling and genotype calling are identical when analyzing the genome of a single individual as inference of a heterozygous or homozygous non-reference genotype would imply the presence of a SNP (Nielsen et al. 2011; Mielczarek and Szyda 2016). However, when it comes to simultaneously analysing multiple samples, an SNP is identified if at least one individual is heterozygous or homozygous non-reference at a genome position (Nielsen et al. 2011; Mielczarek and Szyda 2016). Many empirical and statistical methods have been developed to perform SNP and genotype calling to discover genetic variants accurately. Most of them are based on either heuristic or probabilistic methods.

#### 3.4.1 Heuristic methods

For variant detection, a heuristic algorithm determines genotypes based on a filtering step where variants fulfilling pre-set thresholds are kept. Such thresholds include coverage (e.g. minimum =33), base quality (e.g. minimum = 20) or variant allele frequency (e.g. minimum = 0.08). Then, each allele supported by Fisher’s exact test of read counts is applied compared to the expected distribution based on sequencing error alone (Mielczarek and Szyda 2016). Several software tools are developed based on this methods, such as `VarScan2` (Koboldt et al. 2012). However, these methods can be improved by using more empirically determined cutoffs (Nielsen et al. 2011).

#### 3.4.2 Probabilistic methods

More recent approaches integrate several sources of information within a probabilistic framework. One of the advantages of a probabilistic framework is that it facilitates SNP calling in regions with medium to low coverage compared with heuristic methods (Altmann et al. 2012). For moderate or low sequencing depths, fixed cutoffs based genotype calling will lead to under-calling of heterozygous genotypes and some information will inevitably be lost when using static filtering criteria (Nielsen et al. 2011). Another advantage of probabilistic methods is that it can account for uncertainty in genotype call, making it possible to monitor the accuracy of genotype calling (Altmann et al. 2012; Mielczarek and Szyda 2016). Additional information concerning allele frequencies and/or patterns of linkage disequilibrium (LD) can thus be incorporated in downstream analysis (Nielsen et al. 2011; Mielczarek and Szyda 2016).

SNP and genotype calling based on probabilistic methods usually involve the calculation of genotype likelihood which can be used to evaluate the quality scores for each read. To improve the accuracy of calculation of genotype likelihoods, the following parameters can be considered: 1) a weighting scheme should be used that takes correlated errors into account (Li et al. 2008a), 2) recalibrating the per-base quality scores by using empirical data will also improve genotype likelihoods (Nielsen et al. 2011) and 3) information from the SNP-calling step should be incorporated into the genotype-calling algorithm, leading to genotype likelihoods that are calculated by conditioning on the site containing a polymorphism (Nielsen et al. 2011). By adopting a Bayesian approach for variant calling, prior genotype probabilities can be combined with the estimated genotype likelihood to calculate the posterior probability of a particular genotype. The genotype with the highest posterior probability will then generally be chosen and this probability or the ratio between the highest and the second highest genotype probabilities will be used as a measure of confidence in the variant call (Nielsen et al. 2011; Li et al. 2013; Mielczarek and Szyda 2016).

A prior probability of genotype must be assumed because it is a prerequisite to be able to calculate the posterior probability for a genotype. When data is analysed from a single individual, either a uniform prior probability is chosen that assign equal probability to all genotypes or a non-uniform prior can be based on external information, such as dbSNP (SNP database) entries, the reference sequence or an available population sample (Nielsen et al. 2011; Li et al. 2013). Jointly analyzing multiple individuals will improve the prior probabilities by considering allele or geno-type frequencies, using for example maximum likelihood (Li et al. 2009b; Martin et al. 2010) and then by using a Hardy–Weinberg equilibrium (HWE) assumption or other assumptions that relate allele fre-quencies to genotype frequencies to calculate genotype probabilities (Nielsen et al. 2011). Moreover, a significant improvement in genotype-calling accuracy can be achieved by considering linkage disequilibrium (LD) information (1000 Genomes Project Consortium 2010). However, this approach is not very efficient for rare mutations.

#### 3.4.3 Commonly used software tools based on probabilistic methods

Several software tools have been developed that combine probabilistic framework with Bayesian analysis for variant calling, such as `Samtools` (Li et al. 2009c), `GATK` (DePristo et al. 2011) and `FreeBayes` (Garrison and Martin 2012). All these software also support the use of multiple sample SNP calling. `SAMtools` perform SNP and genotype calling in two steps:1) `mpileup` in `SAMtools` computes the possible genotype likelihood and stores those information in BCF (Binary call formats), and 2) `BCFtools` from the `Samtools` packages applies the prior and does the actual variant calling based on genotype likelihoods information calculated in the previous step (Li et al. 2009c;Wang et al.2015). Compared with `Samtools`, `GATK`, based on similar algorithms, features an advantage of automatically applying several filters before processing variant calling or other pre- and post-processing steps, e.g. filtering out reads that fail quality checks and reads with a mapping quality of zero. `HaplotypeCaller`, the most popular SNP and genotype calling module in `GATK`, can discard the existing mapping information and completely reassembles reads in the region whenever a region show signs of variation. It thus allows for more accurate calling in regions that are traditionally difficult to call, such as regions containing different types of variants close to each other. Four steps will be performed by using `HaplotypeCaller` in `GATK`: 1) determining the regions of the genome that it needs to operate on by detecting significant evidence for variation; 2) determining haplotypes by local assembly of the active region by building a De Bruijn-like graph and by identifying potential variant sites by realigning each haplotype against the reference haplotype, 3) determining likelihoods of the haplotypes given the read data by a pairwise alignment of each read against each haplotype using the `PairHMM` algorithm and 4) assigning sample genotypes based on Bayes’ rule. Liu et al. (2013) compared the performance of four common variant callers, `SAMtools, GATK, glftools` and `Atlas2`, using single and multi-sample variant-calling strategies, and came to the conclusion that `GATK` had several advantages over other variant callers for general purpose NGS analyses. Pirooznia et al. (2014) conducted a series of comparisons between single nucleotide variant calls from NGS data and gold-standard Sanger sequencing in order to evaluate the accuracy of each caller module. They found that `GATK` provided more accurate calls than `SAMtools`, and the `GATK HaplotypeCaller` algorithm outperformed the older `UnifiedGenotype` algorithm (Pirooznia et al. 2014).

Based on these results, we have performed SNP and genotype calling by using `GATK HaplotypeCaller` by generating intermediate genomic VCFs (gVCFs) on a per-subset and per-sample basis (20 gVCFs produced for each individual). These gVCF files were then used for joint calling of multiple samples by using the `GenotypeGVCFs` module in `GATK`. SNP and genotype calling by `GATK` produced 20 VCF files with each file including variants from all 35 individuals (Figure 1). The raw variant calls are likely to contain many false positives arising from errors in the genotyping step or form incorrect alignment of the sequencing data and the called variants therefore needs to be subjected to a number of subsequent filtering steps (e.g. missing data, depth and mapping quality) to produce data that is of sufficient quality for answering the biological questions of interest.

## 4. Variant filtering

### 4.1 Filtering for depth and excessive heterozygosity

A first filtering step was conducted on each of the 20 genomic subset VCF files separately with `vcftools` (Danecek et al. 2011) and `bcftools` (Li et al. 2009) to only include biallelic SNPs positioned > 5 bp away from an intron and where the SNP quality parameters fulfilled `GATK` recommendations for hard filtering (https://gatkforums.broadinstitute.org/gatk/discussion/2806/howto-apply-hard-filters-to-a-call-set).

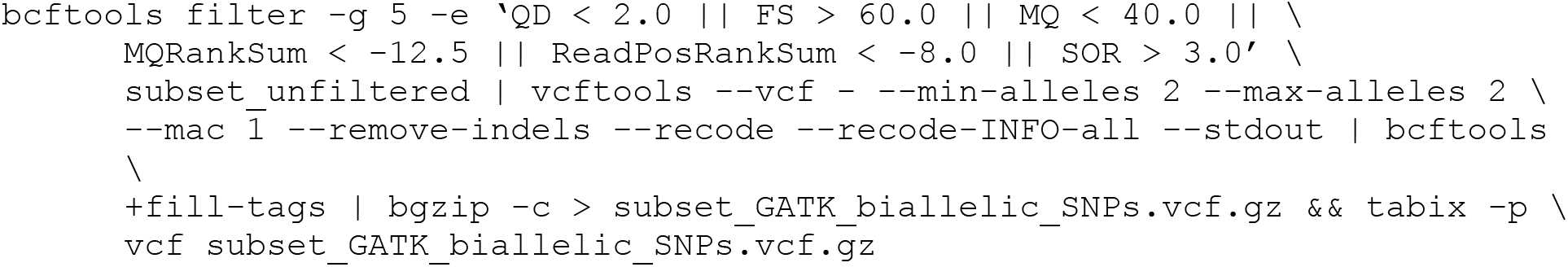

In addition to the usual problems of calling variants from data with low coverage, SNP calling in conifers face the extra problem of possible collapsed repetitive regions in the assembly. The spruce genome, as most conifers, is highly repetitive and the v1.0 *P. abies* assembly is missing approximately 30% of the predicted genome size (Nystedt et al 2013). These genome regions are either completely absent from the reference assembly or present as collapsed regions of high sequence similarity. This introduces problems for read mapping and variant calling as it increase the probability of calling false SNPs in collapsed regions since reads that in reality derive from different genomic regions are mapped at a single region in the assembly. In order to reduce the impact of these issues on variant calling, we performed a second filtering step so that genotype calls with a depth outside the range 6-30 and a GQ < 15 were re-coded to missing data. We then filtered each SNP for being variable with an overall average depth in the range of 8-20 and a ‘maximum missing’ value of 0.8 (max 20% missing data). All info tags were then recalculated with `bcftools fill-tags`.

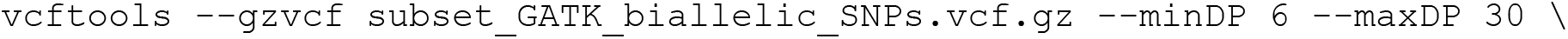
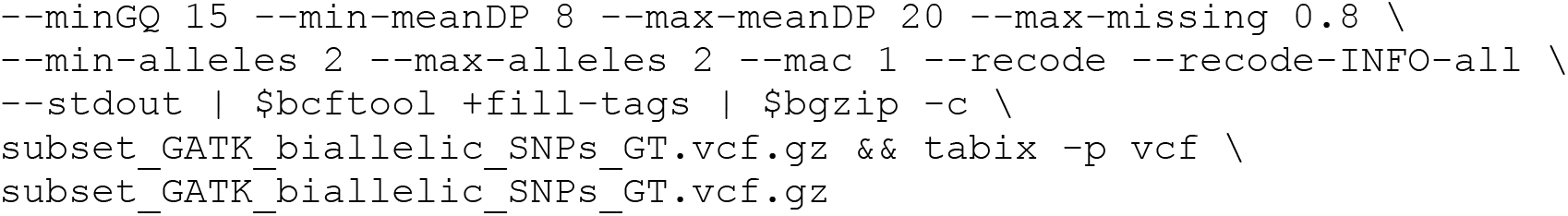

Finally, to remove obviously collapsed SNPs we filtered out all SNPs that displayed a *p*-value of excess of heterozygosity less than 0.05. This was done since SNPs called in collapsed regions should show excess heterozygosities since they are based on reads that are derived from different genomic regions.

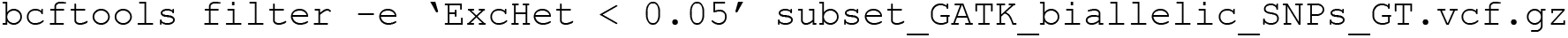

In order to evaluate the filtering parameters, we extracted summary statistics for each of the 20 genomic subsets and filtering steps using `bcftools stats` (Li et al. 2009). We also wrote a custom python script (v2.7) that extracted basic statistics regarding deviations from Hardy-Weinberg equilibrium (`HWE`), excess of heterozygosity (`ExcHet`), number of called samples, allele frequency, number of heterozygous calls, alternative allele ratio for heterozygous calls as well as total depth and total heterozygous depth, from the VCF file of two of the genomic subsets (subset 5 and 6) using pysam (https://github.com/pysam-developers/pysam, Li et al. 2009c). This data was then analysed in R (R Core Team 2014).

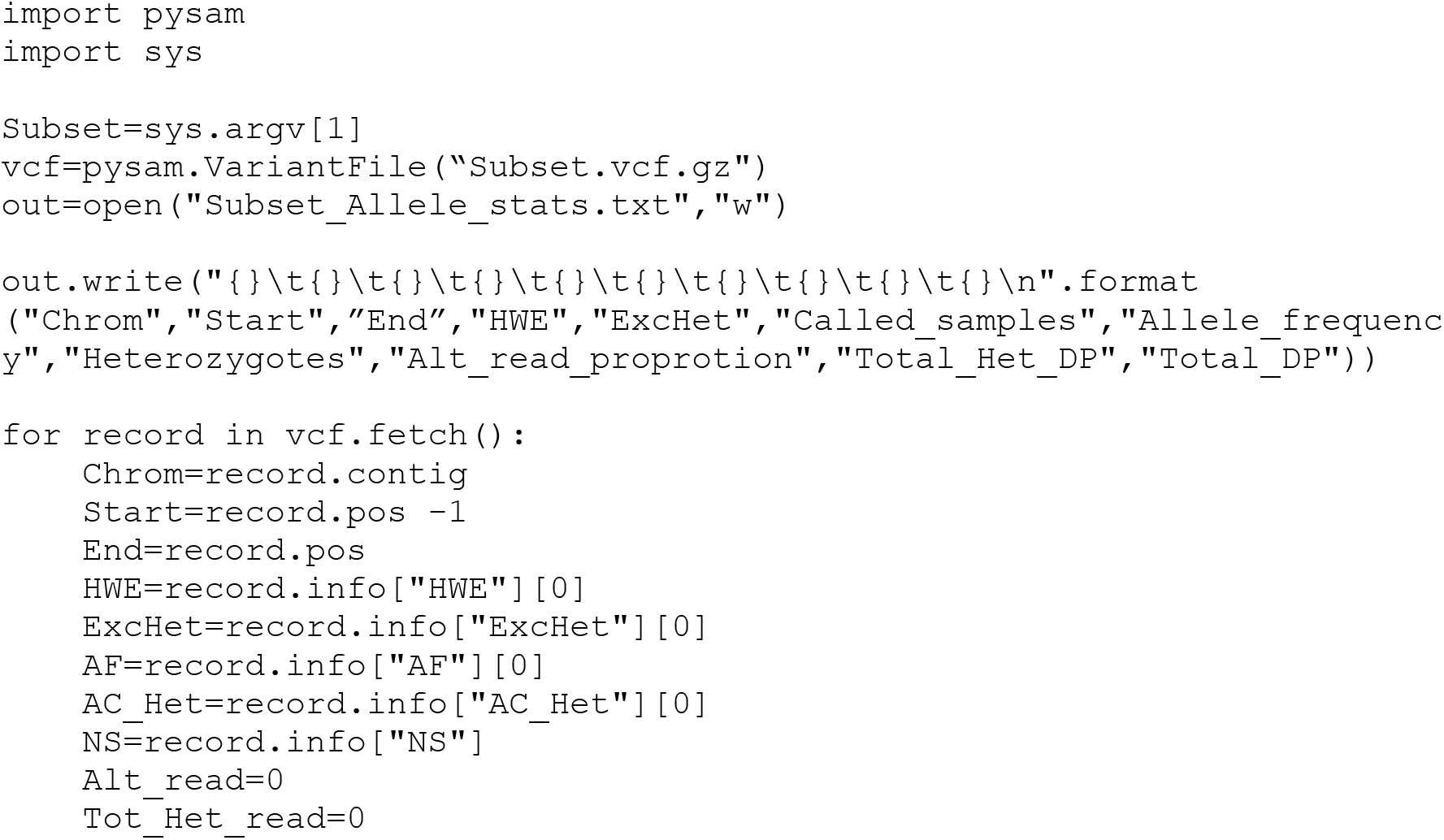
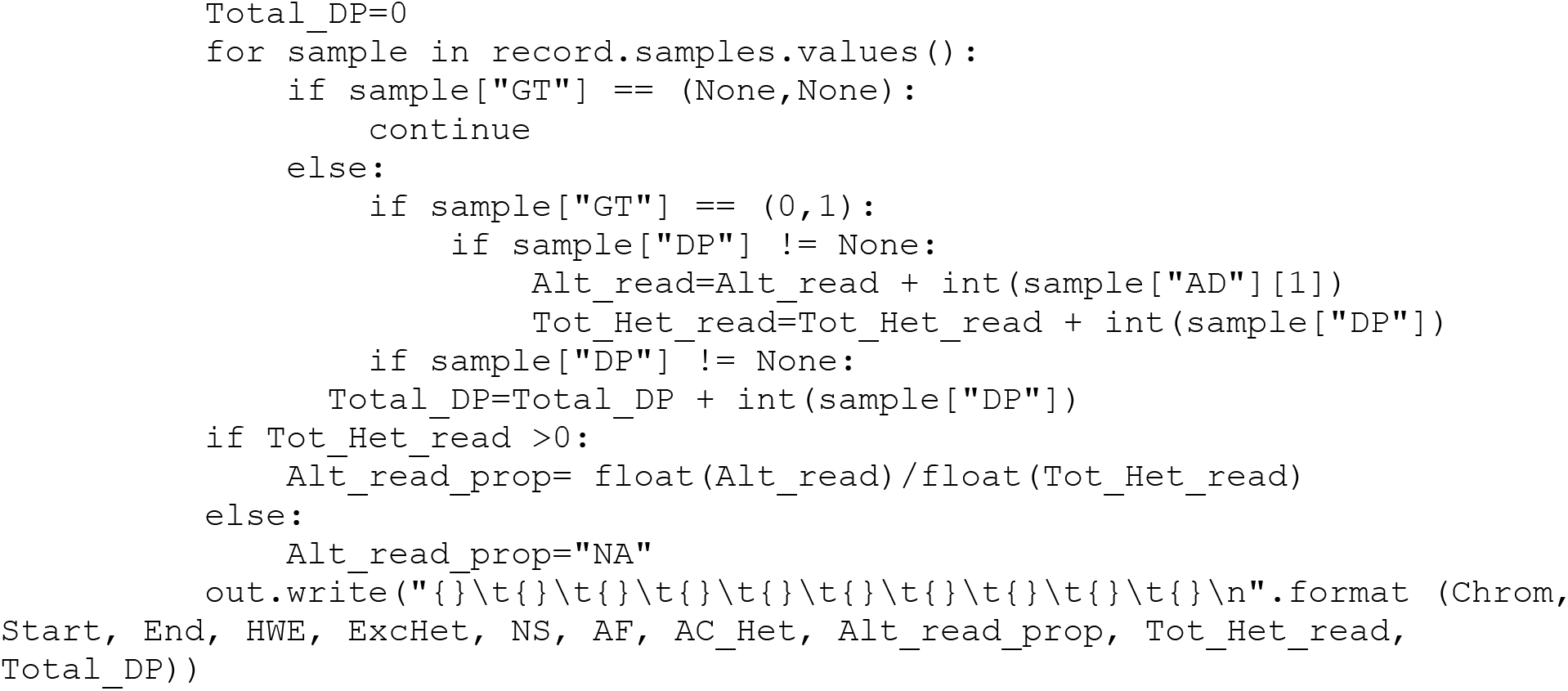

For analyses of different genomic regions separately (ie. repeat regions, outside repeat regions, genic regions and exonic regions), we subset the allele statistics file (above) to corresponding regions in files `Pabies1.0-all.phase.changed.gff3` and `Pabies1.0_Repeats_2.0_repeatmasker.gff3.gz` (available from ftp://plantgenie.org/Data/ConGenIE/) using a custom made python script (v2.7) and `BEDTools` (Quinlan AR and Hall IM, 2010; Dale RK, Pedersen BS, and Quinlan AR. 2011).

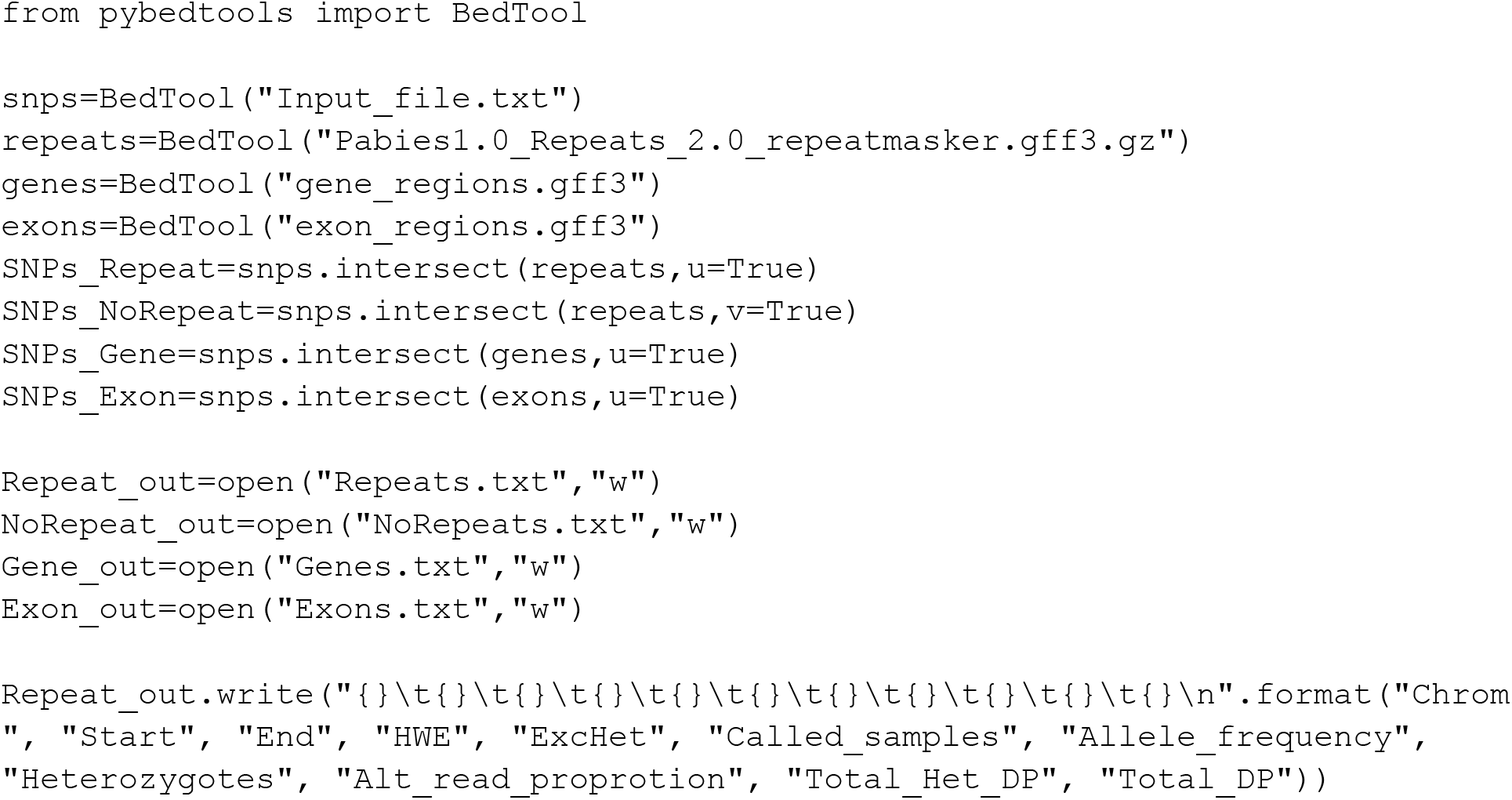
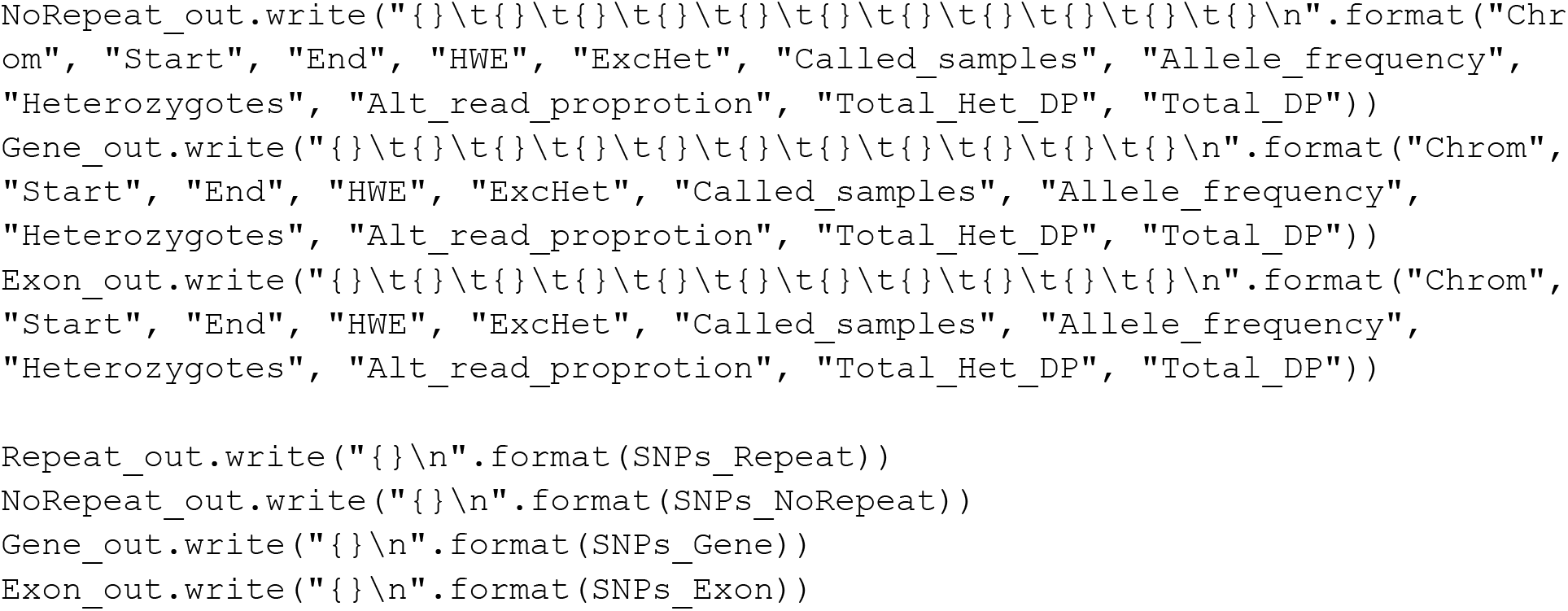

### 4.2 Results

The raw unfiltered VCF files contain a total of 750 million records of which 710 million were single nucleotide polymorphisms (SNPs). After hard-filtering using `GATK` best-practice recommendations (https://gatkforums.broadinstitute.org/gatk/discussion/2806/howto-apply-hard-filters-to-a-call-set), 545 million SNPs (76.8%) remained. These variants were distributed over 94.7% of the 1,970,460 scaffolds in the assembly that are longer than 1 Kb (Table 2). When comparing the alternative allele frequency with the proportion of heterozygous calls per SNP, it is obvious that we have a high proportion of SNPs that do not follow Hardy-Weinberg expectations (Figure 2A). SNPs with either too high or too low fraction of heterozygous calls most often also show a greater deviation in allele ratio in their heterozygous calls compared to SNPs following H-W expectations (Figure 2B-D).

**Figure 2:**
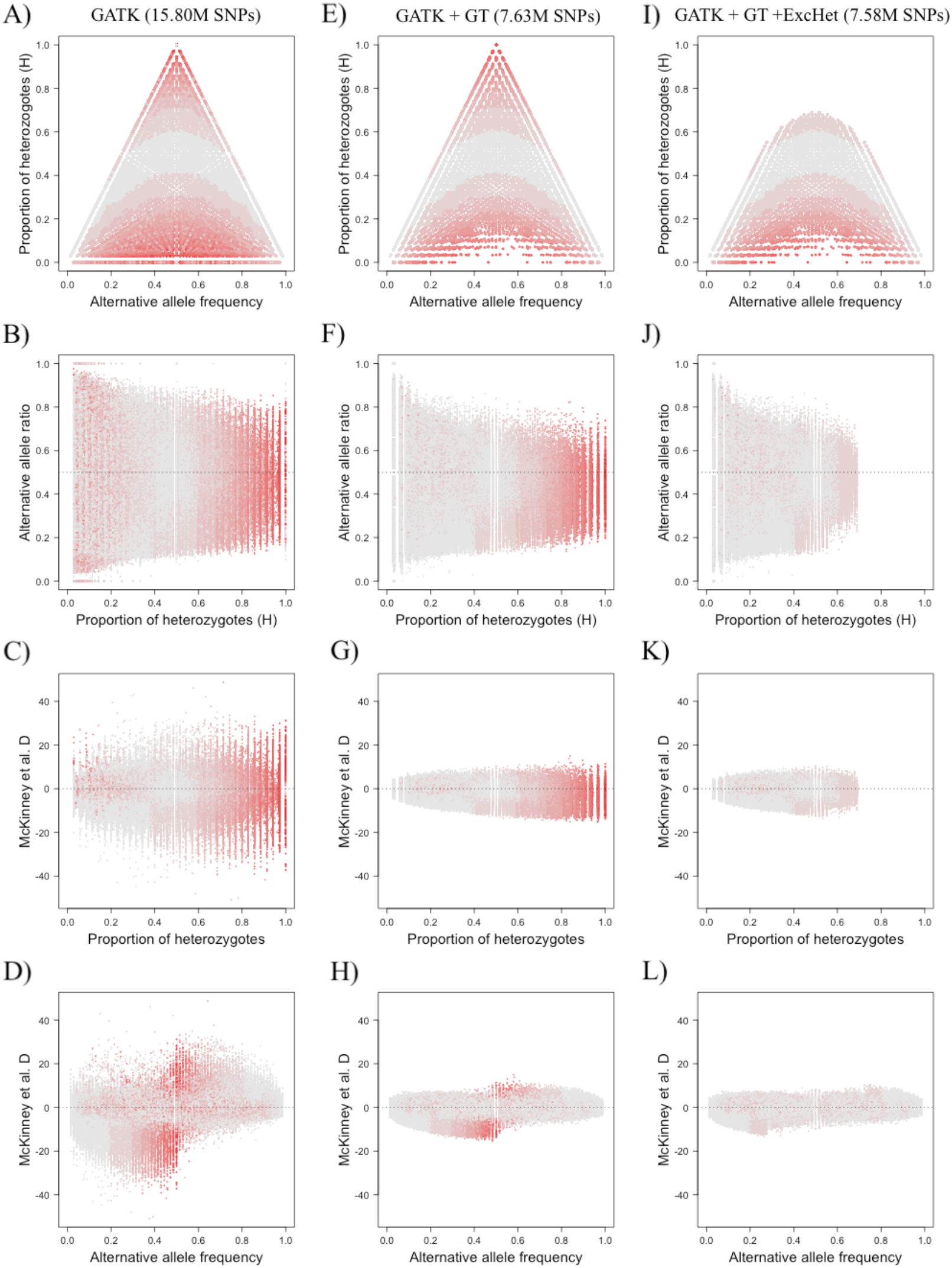
Sub 6 of the WGS data with `GATK` filtered SNPs (A-D, 15.8M SNPs), `GATK + GT` filtered SNPs (E-H, 7.63M SNPs), and `GATK + GT + ExcessHet` filtered SNPs (I-L, 7.58M SNPs) The first row shows the alternative allele frequency versus proportion of heterozygotes for each SNP. The second row shows the proportion of heterozygotes versus alternative allele ratio of all heterozygotes for each SNP. The third row shows the proportion of heterozygotes versus the standardized deviation of allele ratio (McKinney *et al.* 2017) for each SNP. The fourth row shows the alternative allele frequency versus the standardized deviation of allele ratio (McKinney *et al.* 2017) for each SNP. Each SNP is colorised according to Hardy-Weinberg deviations, going from grey for SNPs matching Hardy-Weinberg expectations to bright red for SNPs showing strong deviations.

In genotyping by sequencing (GBS) data, often applied to species that lack a reference genome, duplicated regions have been shown to behave in a very similar manner (McKinney et al. 2017) to our whole genome sequencing (WGS) data. To overcome the issues relating to false SNPs that stem from collapsed/ duplicated genomic regions, McKinney et al. (2017) suggests filtering for sequencing depth, so that all accepted calls have a coverage within the expected span (based on the estimated sequencing depth). We therefore re-coded individual genotypes to missing data if they had a depth that was too low to reliable call both alleles (< 6 reads covering a site), or if their depth was too high compared to the overall coverage (> 30 reads covering a site), which makes it likely that reads are derived from multiple genomic positions. Calls with a genotype quality (GQ) less than 15, which indicates that the difference in likelihood between the best and second best genotype is small, were also re-coded to missing data. We then filtered on an overall average depth per SNP of 8-20 to reduce the proportion of possible erroneous calls further and also added a threshold of maximum 20% missing data in order to keep a SNP for downstream analysis.

296 million SNPs, corresponding to 41.7% of the raw unfiltered SNPs, remained after these filtering steps and were distributed over 63.2% of the >1Kb scaffolds (Table 2). When comparing the proportion of heterozygous calls to the alternative allele frequency, a noticeable amount of the SNPs with heterozygous deficiency has been filtered out (Figure 2E). The deviation from a balanced allele ratio of heterozygous calls is also visibly lower (Figure 2F-H) even though the data still suffers from SNPs showing heterozygous excess at intermediate alternative allele frequencies (Figure 2H at ~0.5). Such excess of heterozygous could be explained by real biological phenomena, such as balancing selection (Charlesworth 2006), or arise due to artefacts caused by collapsed/duplicated regions in the assembly as discussed above (McKinney et al. 2017). Regions under balancing selection should be highly localised in the genome while the risk of collapsed regions in the spruce assembly is *a priori* expected to be high since we know that the genome contains a huge proportion of retrotransposon-derived sequences (Nystedt et al. 2013, Cossu et al 2017). To filter out SNPs with a significant excess of heterozygotes therefore seems justifiable for the data at hand. Heterozygous deficiency, on the other hand, can be both explained by errors in variant calling in low-depth regions or by population structure (Hartl and Clark 1989). The latter is a much more likely scenario for our data set, since samples are derived from all over the distribution range of *P. abies* and we also expect this pattern to manifest itself across most of the genome. We therefore chose not to filter on this criteria. By removing all SNPs showing a significant excess of heterozygotes (*p*-value < 0.05), we ultimately retained 294 million SNPs (corresponding to 41.4% of the unfiltered SNPs) from 63.2% of the > 1kb scaffolds (Table 2, Figure 2I-M).

### 4.3 Comparison of WGS and reduced representation sequencing data

Even with a number of conifer draft genomes available (Nystedt et al. 2013, Stevens et al 2016, Zimin et al 2017, Neale et al 2017), the method of choice for analysing sequence diversity in conifers has been different kinds of reduced representation sequencing methods, such as genotyping by sequencing (GBS), restriction site associated DNA sequencing (RadSeq) or capture probe sequencing (Syvänen 2005, Andrews et al. 2016). These methods all have in common that they target a small fraction of the target genome, the only thing that differ is how this fraction is selected. In order to compare the behaviour of our spruce WGS data with data derived from a reduced representation sequencing technique, we analysed at set of 526 samples that had been genotyped using a set of 40,018 sequence capture probes that had been designed to target regions within genic regions of the v1.1 *P. abies* genome assembly (Baison et al. 2018, Vidalis et al. 2018). The same filtering steps as described in the preceding section were used for the capture probe data set, although the thresholds for individual depth range (6-40), overall average depth range (10-30) and significance level for excess of heterozygotes (*p*-value <1e-10) were altered to fit the size of data set (the number of samples and estimated sequencing depth). Even though the probes were designed to target unique regions in the assembly (Vidalis et al. 2018), the `GATK` SNP quality filter show a remarkably high level of SNPs with deviations from Hardy-Weinberg equilibrium (Figure 3A) with large deviations from balanced allele ratios in heterozygous calls (Figure 3B-D).

**Figure 3:**
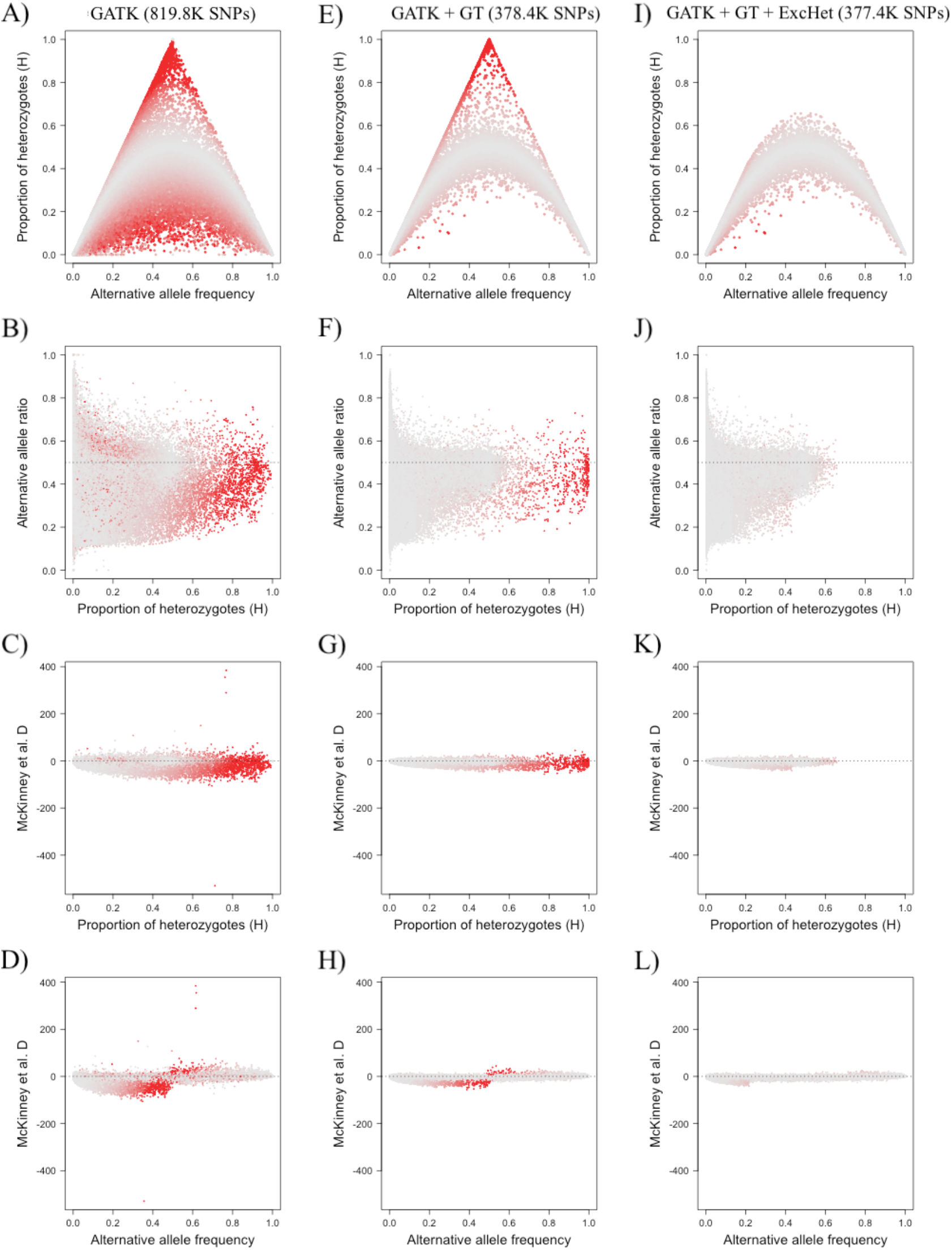
Sequence capture data of 526 individuals with `GATK` filtered SNPs (A-D, 819.8K SNPs), `GATK + GT` filtered SNPs (E-H, 378.4K SNPs), and `GATK + GT + ExcessHet` filtered SNPs (I-L, 377.4K SNPs) The first row shows the alternative allele frequency versus proportion of heterozygotes for each SNP. The second row shows the proportion of heterozygotes versus alternative allele ratio of all heterozygotes for each SNP. The third row shows the proportion of heterozygotes versus the standardized deviation of allele ratio (Mackinnely et al …) for each SNP. The fourth row shows the alternative allele frequency versus the standardized deviation of allele ratio (Mackinnely et al …) for each SNP. Each SNP is colorised according to Hardy-Weinberg deviations, going from grey for SNPs matching Hardy-Weinberg expectations to bright red for SNPs showing strong deviations.

Adding the depth filters provides a huge improvement in SNP quality by both reducing the amount of SNPs showing excess of heterozygosity as well as improving the balance in allele ratios for heterozygous calls. The sequence capture data set does not show the same level of heterozygote deficiency as wee seen in the WGS data (Figure 2 vs Figure 3). However, the capture probe data set only contain plus trees sampled from Southern/Central Sweden (Baison et al. 2018) and it is therefore not surprising we observe less effects of population structure in this data, since these trees cover a much smaller geographic region than the samples used in the WGS data. Nevertheless, even after removing SNPs showing an excess of heterozygotes, we see a bias towards the reference allele in the sequence capture data that is not visible in the WGS data (bias towards allele frequencies <0.5 in Figure 3J). This is most probably due to an artefact of using probes, since the probes were designed against the reference alleles of the genome and likely work better in regions with low to moderate amounts of variation (Vidalis et al. 2018).

To evaluate the proportion of collapsed regions within genic regions even further, we analysed 1997 haploid sequence captured samples that had previously been mapped to the whole genome, called using a diploid ploidy setting and subset to the probe regions ± 100 bp (Bernhardsson et al. 2018, Vidalis et al. 2018). These samples, which should only have homozygous calls, showed an average heterozygosity level of approximately 3.7% (obviously contaminated samples with heterozygosity levels > 10% removed). 114K SNPs remained after `GATK` hard filtering and with no more than 30% missing data, and these showed a median heterozygosity level of 0.01 (1st and 3rd quartile of 0.004 and 0.017, respectively). However, 10% of the SNPs experienced a heterozygosity level > 0.05 (corresponding to 70-98 heterozygous calls depending on call rate), with a maximum of 0.95 (data not shown).

### 4.4 Comparisons of genic, inter-genic and repetitive regions

In order to analyse how different regions of the spruce genome behave with regards to deviations from Hardy-Weinberg equilibrium, we subdivided the `GATK` SNP quality + depth and genotype quality filtered WGS data set into four sets: inside repeat regions (regions covered by known repetitive elements), outside repeat regions (regions outside known repetitive elements), genic regions (all regions falling within an annotated gene model) and exonic regions (all regions falling within exons of annotated gene models). We then applied the same analysis as described earlier to assess the proportion of heterozygous calls versus alternative allele frequency and alternative allele ratios to each of the four sets. Interestingly enough, both the ‘within repeat regions’ and ‘outside repeat regions’ data sets behave similarly, showing both an excess and a deficiency of heterozygous calls (Figure 4A-H). Genic and exonic regions, on the other hand, show much fewer SNPs that deviate from Hardy-Weinberg expectations with an overall pattern much more similar to what we observed in the sequence capture data set (Figure 4I-P).

**Figure 4:**
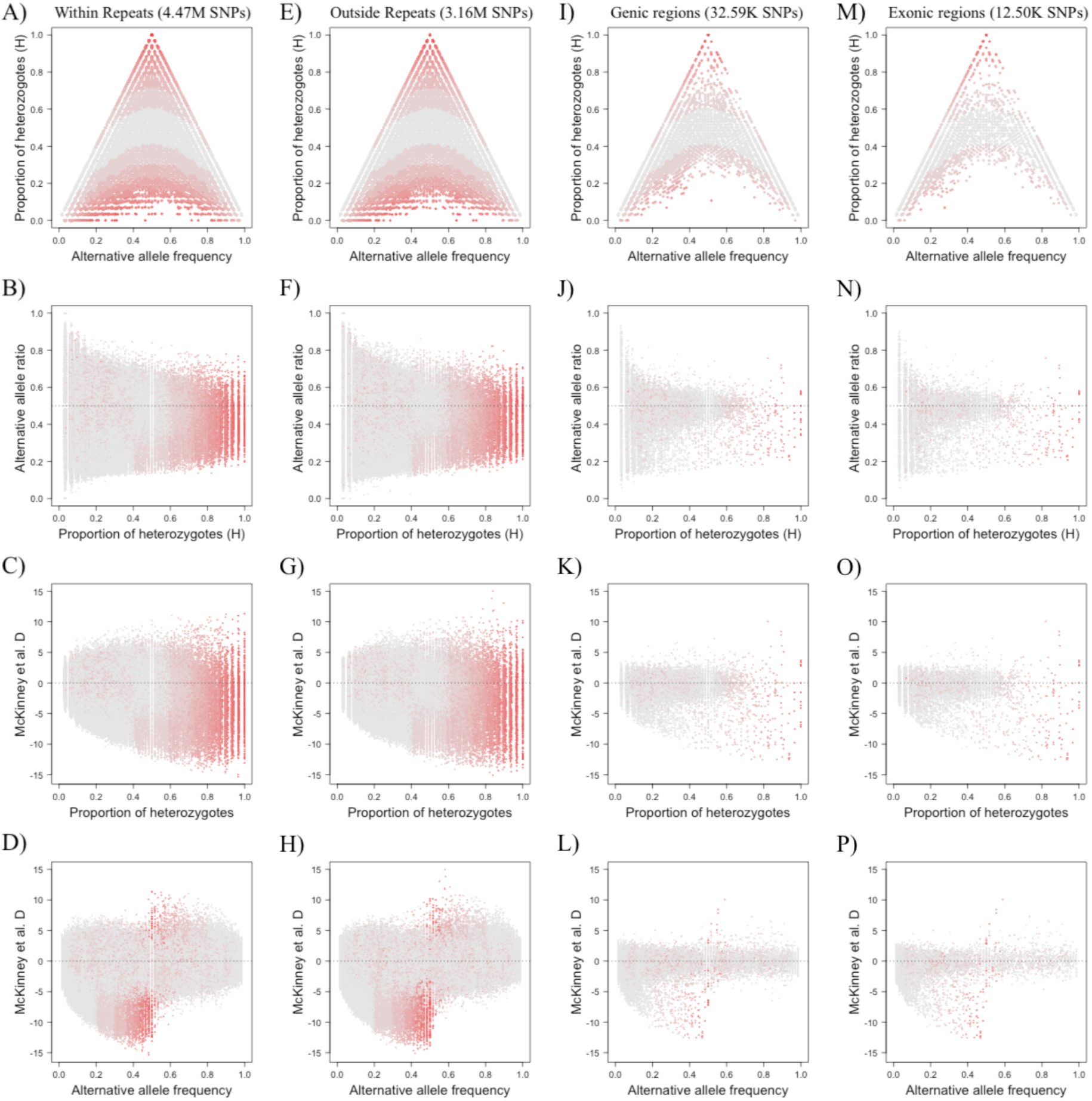
Subset 6 of the WGS data with `GATK + GT` filtered SNPs divided into four regions. SNPs within repeat regions (A-D, 4.47M SNPs), SNPs outside repeat regions (E-H, 3.18M SNPs), SNPs within genic regions (I-L, 32.6k SNPs) and SNPs within exonic regions (M-P, 12.5k SNPs). The first row shows the alternative allele frequency versus proportion of heterozygotes for each SNP. The second row shows the proportion of heterozygotes versus alternative allele ratio of all heterozygotes for each SNP. The third row shows the proportion of heterozygotes versus the standardized deviation of allele ratio (Mackinnely et al …) for each SNP. The fourth row shows the alternative allele frequency versus the standardized deviation of allele ratio (Mackinnely et al …) for each SNP. Each SNP is colorised according to Hardy-Weinberg deviations, going from grey for SNPs matching Hardy-Weinberg expectations to bright red for SNPs showing strong deviations.

In order to understand how the two filtering steps, `GATK` best practice SNP quality filters (hereafter called `GATK`) and `GATK` practice SNP quality filters and depth and genotype quality (hereafter called `GATK+GT`), change the SNPs retained across the different genomic subsets, we analysed summary statistics from all subsets. The fraction of SNPs retained in subsest after the `GATK` filter were strongly negatively correlated with fraction of sites covered by repeats in the subsets (Figure 5A, correlation −0.92,p-value=1.4e-8). This correlation was reduced by using the `GATK+GT` filtering criteria to the subsets (Figure 5A, correlation between fraction retained SNPs and repeat coverage of −0.77, p-value=8.1e-5), but remained negative resulting in fewer SNPs being called in genomic subsets containing higher fraction of repetitive sequences. The median physical distance between SNPs increased from 16.5 bp for the `GATK` filtered subsets (ranging from 15.0 to 55.7 bp per subset, with an overall average of 17.3 bp) to 36.3 bp for the `GATK+GT` filtered subsets (ranging from 23.2 to 512.8 bp per subset with an overall average of 32.0 bp) (Figure 5C, outliers not shown). Even though this suggest a high level of nucleotide diversity in Norway spruce, most alleles are rare and as many as 26% of the SNPs appear as singletons in the `GATK+GT` filtered data set, compared to 21% in the `GATK` filtered data set. The `GATK+GT` filter has a large impact on the fraction of scaffolds that retain any SNPs, decreasing from 94.7% for the `GATK` filtered data set to only 63.2% under `GATK+GT` (Figure 5B). The transition-transversion ratio similarly decreased with the `GATK+GT` filter (Figure 5D).

**Figure 5:**
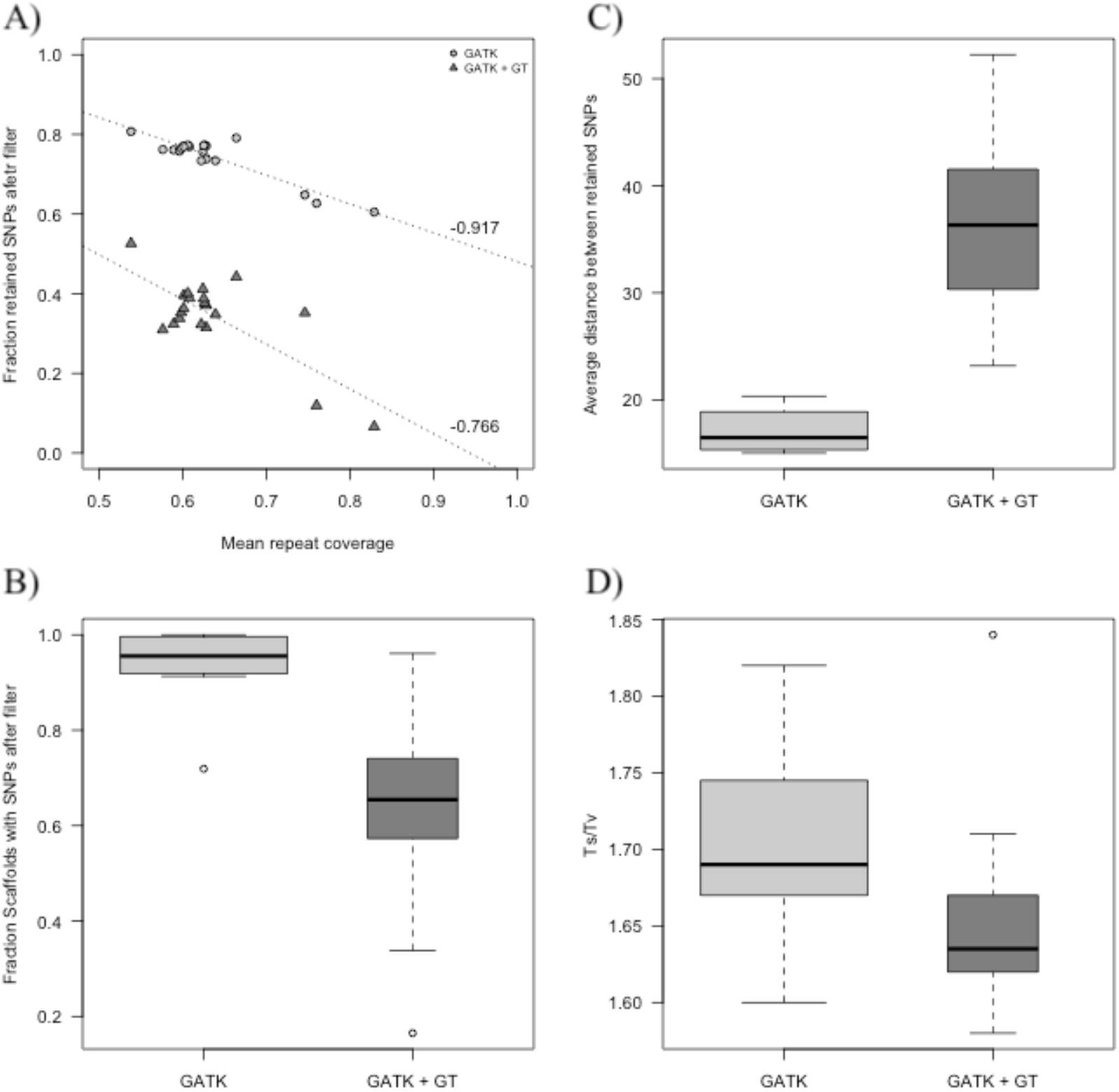
Summary statistics on a per genomic subset basis, over all samples. GATK indicate the GATK SNP quality filtered data set; GATK+GT indicate the GATK SNP quality + depth and genotype quality filtered data set. A) Correlation between proportion of retained SNPs after filter (in comparison to the amount of unfiltered raw SNPs) and the average level of repeat coverage. B) Proportion of scaffolds with retained SNPs after filter. C) Average physical distance (in bp) between retained SNPs. outliers are not shown. D) Transition-transversion ratio. The width of the boxes are proportional to the fraction of retained SNPs after filter.

At the individual sample level, the `GATK+GT` filter resulted in an increased proportion of heterozygous calls and a reduced proportion of homozygous alternative calls, while the proportion of homozygous reference calls stayed the same in comparison to the `GATK` filter (Figure 6A-B,D). Sample *Pab006*, which is the individual from which the *P. abies* reference assembly was derived (Z4006, Nystedt et al. 2013), have SNPs called as an homozygous alternative, a genotype which should be impossible in this individual, in 0.07% of the calls using the `GATK` filtering criteria. This fraction was reduced to 0.02% when using the `GATK+GT` filtering criteria, suggesting that the depth filter do improve the quality of the genotype calls and gives an overall higher average depth in comparison to the `GATK` filtered data set (Figure 6C). Although the average sequencing depth of called genotypes increase following `GATK+GT` filtering, the proportion of missing calls also increase, and reaches 30% for samples with the lowest estimated sequencing coverage (Figure 6E). The increase in missingness do not affect the proportion of singletons per sample, which stays roughly constant under both the `GATK` (average of 0.8%) as well as for the `GATK+GT` (average of 0.9%) filtering criteria (Figure 6F).

**Figure 6:**
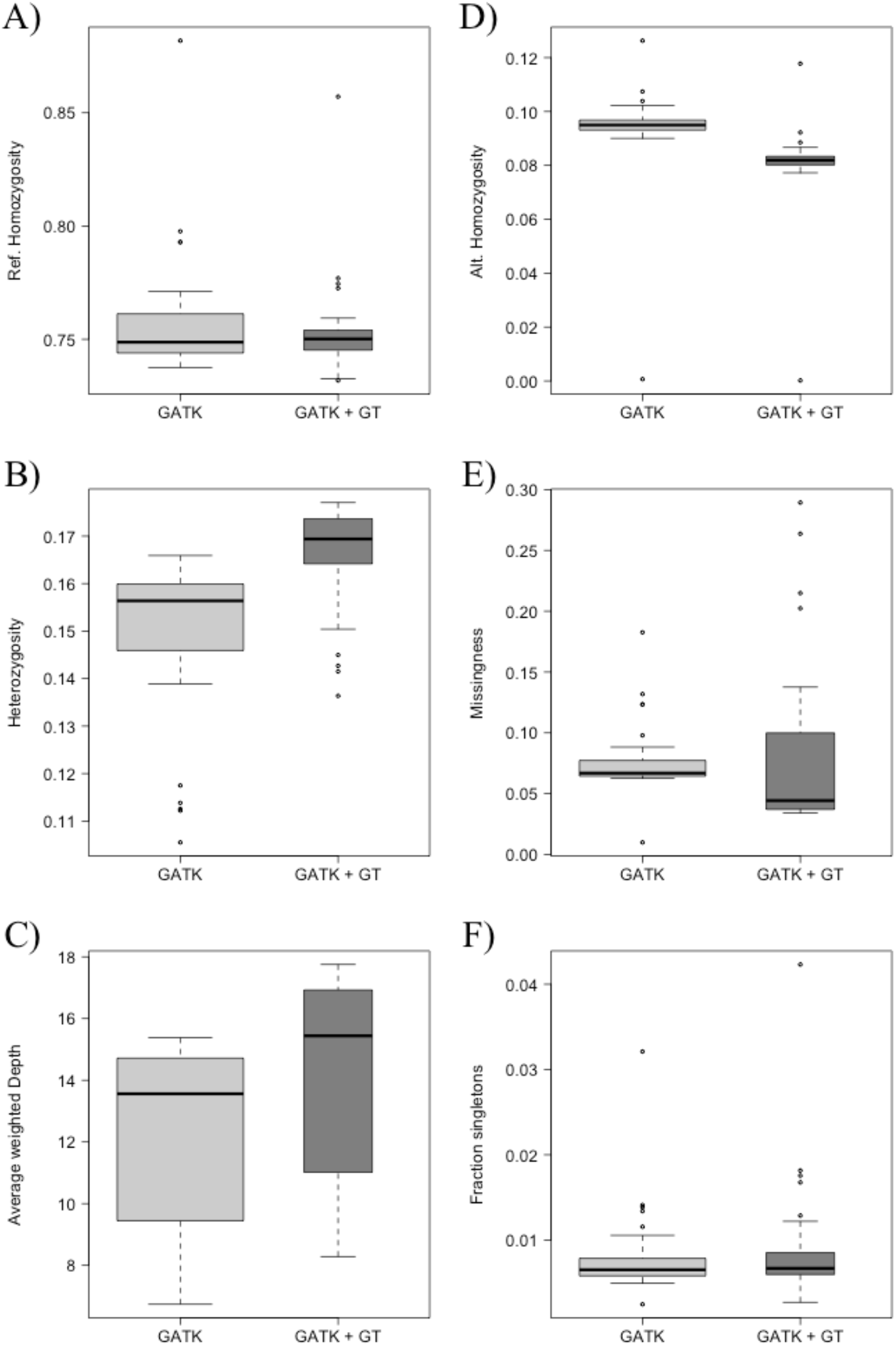
Summary statistics on a per individual basis, over all genomic subsets. GATK indicate the GATK SNP quality filtered data set. GATK+GT indicate the GATK SNP quality + depth and genotype quality filtered data set. A) Proportion of calls that are homozygous reference. B) Proportion of calls that are heterozygous. C) Average depth over all calls. Average depth per genomic subset was weighted towards its number of SNPs before an average was calculated over all genomic subsets. D) Proportion of calls that are homozygous alternative. E) Proportion of SNPs with missing data (No calls). F) Proportion of calls that are singletons. The width of the boxes are proportional to the fraction of retained SNPs after filter.

### 4.5 Effects of filtering on estimates of the site frequency spectrum

To analyse how different summary statistics regarding the site frequency spectrum (SFS) is affected by filtering parameters (`GATK` and `GATK + GT + ExcHet`), we analysed Tajima’s D (Tajima 1989) and pairwise nucleotide diversity (π, Hartl and Clark 1989) on a per scaffold basis for genomic subset 6, for inside repeat regions, outside repeat regions, genic regions and exonic regions separately. The `GATK` filtered data showed an overall higher estimate of Tajima’s D than the `GATK + GT + ExcHet` filtered data for all four genomic regions (average of −0.35 – −0.13 and −0.57 – −0.53 for `GATK` and `GATK + GT + ExcHet`, respectively). These estimates were however highly correlated between the filtering parameters in all four genomic regions (correlation of 0.77-0.82 with p-value < 2.2e-16 for all four genomic regions, Figure 7A-D). The `GATK` filtered data also show an overall slightly higher nucleotide diversity level (π) compared to the `GATK + GT + ExcHet` filtered data (average of 3.7e-4 – 1.6e-3 and 2.1e-4 – 1.0e-3 for `GATK` and `GATK + GT + ExcHet`, respectively), with lower diversity level for the same amount of analysed variants in the fully filtered data set (Figure 7E-H). Smaller differences regarding the SFS between genomic regions could be found in the fully filtered data in comparison to the `GATK` filtered data, indicating that we managed to remove false SNPs due to collapsed genomic regions in the assembly without altering the SFS by filter for minor allele frequencies.

**Figure 7:**
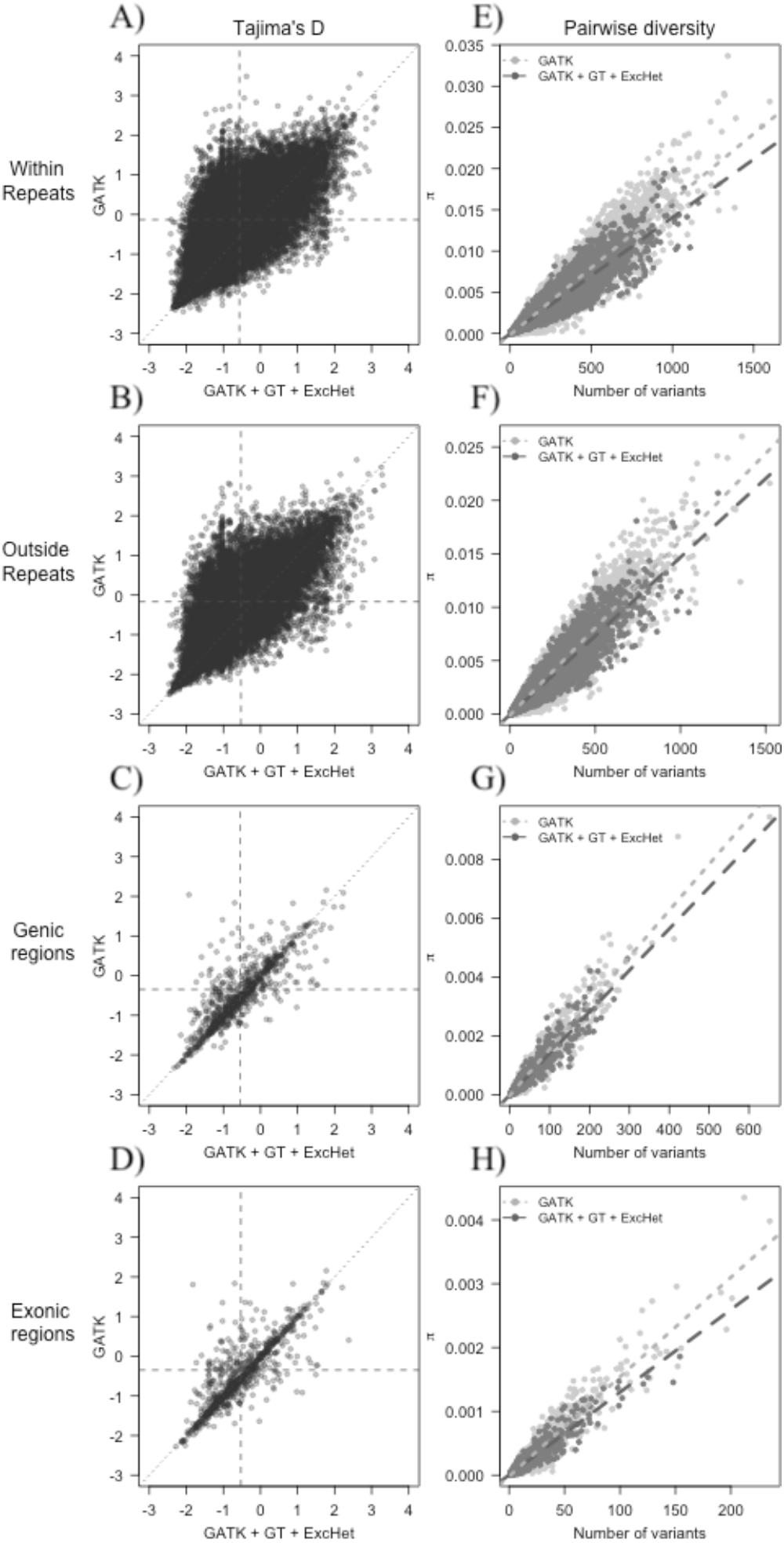
Analyses of the site frequency spectrum (SFS) for genomic subset 6, divided into within repeat regions (row 1), Outside repeat regions (row 2), genic regions (row 3) and exonic regions (row 4). A-D) Correlation between Tajima’s D estimated over scaffolds for the `GATK` and `GATK + GT+ExcHet` filtered data sets. Horizontal dashed line indicate average Tajima’s D för the `GATK` filtered data. Vertical dashed line indicate average Tajima’s D för the `GATK + GT + ExcHet` filtered data. Diagonal dotted line shows the 1:1 correlation between the data sets. E-H) Estimates of pairwise nucleotide diversity (π) per scaffold versus number of analysed variants for `GATK` filtered data (light grey dots and dotted line) and `GATK + GT + ExcHet` filtered data (dark grey dots and dashed line). Lines shows the linear model (lm) between diversity and number of analysed variants for each of the data sets.

## 5. Conclusions and recommendations

Having high-quality variant calls is essential for downstream analyses such as population genomic studies, inferences of demographic history or complex trait dissection using, for instance, genome wide association studies (Nielsen et al 2011). Variant calling in conifers pose a number of challenges arising from the complex highly repetitive nature of conifer genomes. Even in the most complete reference genomes of conifers, large regions of the genome are still lacking and this cause problems for mapping of short (<250bp) sequencing reads, since reads from such regions will not match any region in the reference genome. Such reads may fail to map altogether and will be eliminated from further steps in the variant calling pipeline. However, if they are derived from repetitive regions, highly similar, genomic regions can be represented in the assembly allowing read mapping. To a lesser degree these issues are true also for sequencing reads from repetitive regions that are represented in the assembly as these reads may match equally well to multiple regions in the reference genome. The end result in both cases is thus that some regions are being covered by sequencing reads originating from different genomic regions which may ultimately result in the calling of false genetic variants as nucleotide substitutions that differentiate the paralogous regions are called as true SNPs (Gayral et al., 2013). Such variants often can have high quality scores, making it hard to filter them out using even the best practice procedures for variant filtering (Nielsen et al 2011).

These issues are apparent in our data even when using what is considered rather stringent criteria for variant filtering (Figure 2). To alleviate these issues we have implemented further filtering on sequencing depth and on excessive heterozygosity. Filtering on too high depth is expected to target primarily false variants that are derived from possible repetitive regions where as filtering on too low depth targets variants where the low sequencing depth make calling of both alleles in heterozygous individuals unlikely. Filtering on low depth should thus target called variants that may have deflated levels of heterozygosity where as filtering on high depth primarily should target variants with excessive heterozygosity. However, even very stringent filtering on high depth does not completely eliminate problems the excess heterozygosity and we therefore have included a final filtering step aimed specifically at excess heterozygosity (Figure 2).

To a certain extent, having access to better and more contiguous reference genomes will alleviate some of these issues but it will likely not eliminate them completely. So even as conifer reference genomes improve due to, for instance, the use of long-read sequencing technologies re-sequencing and variant calling from short read data will still be difficult and fraught with both false positive and false negative SNPs. One possible solution to these issues is to base re-sequencing on haploid tissue, e.g from megagametophytes. Using haploid tissue has the benefit that heterozygous genotypes should be absent and any heterozygous genotype call observed is thus likely to arise to to either sequencing errors or from mapping of paralogous reads to a single region in the reference genome. Utilizing haploid tissues to rule out issues arising from paralogy has already been used in conifers to improve, for instance, transcriptome assembly (Ojeda et al. 2018). Another possible way forward would be to utilize longer reads for re-sequencing, thereby reducing the probability that a single read may map to multiple regions in the reference genome. Although the cost and throughput of current long-read technologies make genome re-sequencing of conifers using exclusively long reads infeasible, declining costs and technology development could open up possibilities for re-sequencing using long reads in the future.

## 6. Acknowledgements

The research has been funded by grants from the Knut and Alice Wallenberg foundation (Norway spruce genome project) and the Swedish Foundation for Strategic Research (SSF, grant number RBP14–0040). Data generation was supported by Science for Life Laboratory and the National Genomics Infrastructure (NGI) which provided access to massive parallel sequencing. All analyses were performed on resources provided by the Swedish National Infrastructure for Computing (SNIC) at Uppsala Multidisciplinary Centre for Advanced Computational Science (UPPMAX) under the projects b2012141, SNIC 2017/1-438, SNIC 2018/3-529 and uppstore2017066. X.W. was supported by a scholarship from the Chinese Scholarship Council (CSC).

## Notes

### Competing Interest Statement

The authors have declared no competing interest.

